# Evidence for prescribed NK cell Ly49 developmental pathways in mice

**DOI:** 10.1101/2020.05.23.112391

**Authors:** Alberto J. Millan, Bryan A. Hom, Jeremy B. Libang, Suzanne Sindi, Jennifer O. Manilay

## Abstract

Previous studies of NK cell inhibitory Ly49 receptors suggested their expression is stochastic. However, relatively few studies have examined this stochasticity in conjunction with activating Ly49 receptors. We hypothesized that the expression of activating Ly49 receptors is not stochastic and is influenced by inhibitory Ly49 receptors. We analyzed NK cell “clusters” defined by combinatorial expression of activating (Ly49H, Ly49D) and inhibitory (Ly49I, Ly49G2) receptors in C57BL/6 mice. Using the product rule to evaluate the interdependencies of the Ly49 receptors, we found evidence for a tightly regulated expression at the immature NK cell stage, with the highest interdependencies between clusters that express at least one activating receptor. Further analysis demonstrated that certain NK clusters predominated at the immature (CD27+CD11b−), transitional (CD27+CD11b+) and mature (CD27−CD11b−) NK cell stages. Using parallel in vitro culture and in vivo transplantation of sorted NK clusters, we discovered non-random upregulation of Ly49 receptors, suggesting that prescribed pathways of NK cluster differentiation exist. Our data infer that upregulation of Ly49I is an important step in NK cell maturation. Ki-67 expression and cell counts confirmed that immature NK cells proliferate more than mature NK cells. We found that MHC-I is particularly important for regulation of Ly49D and Ly49G2, even though no known MHC-I ligand for these receptors is present in B6 mice. Our data indicate that the regulatory systems controlling the expression of both activating and inhibitory Ly49 receptors are non-stochastic and support the idea that NK cell clusters develop in a non-random process correlated to their maturation stage.

## Introduction

Natural killer (NK) cells are innate lymphocytes which function in immune cell surveillance by the recognition and elimination of cellular targets. Unlike their T and B lymphocyte counterparts, which acquire antigen-specific diversity through genetic recombination events, NK cells generate germline-encoded receptors that can recognize and lyse cellular targets by releasing perforin, granzymes, and secrete regulatory and proinflammatory cytokines. Activating and inhibitory Ly49 receptors employ NK cell signaling pathways, which dictate NK cell effector functions through cytoplasmic immunoreceptor tyrosine-based inhibitory motifs (ITIMs) and immunoreceptor tyrosine-based activating motifs (ITAM) (1–4). NK cell subsets are thought to utilize a “rheostat” process in which NK cell subsets are tuned quantitatively by self-MHC class I ligands corresponding to specific Ly49 inhibitory receptors, which in turn provides a diversity of responsiveness towards cellular targets (5, 6). Through this mechanism, NK cells distinguish healthy from unhealthy cells. However, what regulates the acquisition of specific NK cell Ly49 receptors during NK cell development and maturation is still an unanswered and complex question. In addition, how models derived from the biology of Ly49 inhibitory receptors pertain to the acquisition of Ly49 activating receptors is unresolved.

NK cell subsets that express single as well as overlapping Ly49 activating and inhibitory receptors exist, which may reflect the complexity required to ensure host immunity while maintaining self-tolerance (7). Expression of Ly49 receptors, encoded by the polymorphic and polygenic *Klra* genes located on mouse chromosome 6, is often described as stochastic (8, 9). However, Ly49 receptor acquisition may not be entirely stochastic, as it has been shown that inhibitory Ly49 receptors can be regulated by self-MHC class I (MHC-I) expression and controlled by the Ly49 bidirectional transcriptional regulation of Pro1 and Pro2 (10, 11). In contrast, activating Ly49 receptors lack a defined Pro1 region. Mathematical methods to test the interdependence of expression of individual Ly49 inhibitory receptors on the expression of other Ly49 members in MHC-I-deficient and MHC-I-sufficient (wild type) mice support that Ly49 inhibitory receptor expression may not be independently distributed and thus may not be entirely random (8, 12–15). Previous studies provide support that Ly49H and Ly49D are distinctly influenced by co-expression with inhibitory receptor Ly49C (16) and non-stochastic (10, 17) regulation of the expression of Ly49 activating receptors.

In this study, we tested the hypothesis that expression of Ly49 activating receptors is not stochastic and is influenced by Ly49 inhibitory receptors. To test this hypothesis, we utilized a combination of statistical, in vitro and in vivo approaches. We provide evidence for a prescribed pathway of NK “cluster transitions” in vitro and in vivo, which suggest that Ly49 activating receptor acquisition is directed. Even though no known MHC-I ligand for Ly49G2, Ly49H and Ly49D is known in B6 mice, NK cell cluster distribution is altered in MHC-I deficient mice. Taken together, our data support the idea that NK cell clusters develop in a non-random manner and provide additional evidence that the regulatory system that controls the expression of both Ly49 activating and inhibitory receptors is non-stochastic. Our findings lead to an expanded model of NK cell receptor acquisition during NK development and maturation.

## Materials and Methods

### Mice

C57B6/J, B6.SJL-Ptprca Pepcb/BoyJ, and B6.129P2-B2mtm1Unc/J *β*-2 microglobulin knockout (*β*_2_m^−/−^ KO) mice were obtained from The Jackson Laboratory and bred at the University of California, Merced. Mice of both sexes between the ages of 10 and 28 weeks were used for each experiment. We observed no significant differences between mice of both sexes or different ages, except for one specific study on cell proliferation (please see Results). Mice were housed in specific pathogen-free cages with autoclaved feed. Mice were euthanized by carbon dioxide asphyxiation followed by cervical dislocation. All animal procedures were approved by the UC Merced Institutional Animal Care and Use Committee.

### Flow cytometry (FACS)

Splenic cells were harvested, processed and stained for flow cytometric analysis (FACS) as described (18). Cells were stained with the following antibodies, purchased from eBioscience, Biolegend, Miltenyi Biotec, and BD Biosciences: PE/Cy5-CD3 (145-2C11), PE/Cy5-CD4 (RM4-5), PE/Cy5-CD8a (53-67), PE/Cy5-Gr1 (RB6-8C5), PE/Cy5-CD19 (6D5), PE-CD27 (LG3A10), BV421-NK 1.1 (PK136), BV510-CD11b (M1/70), biotinylated or APC-Ly49D (4E5), APC-Cy7-Ly49G2 (4D11), FITC- or APC-Ly49H (3D10), APC-VID770-Ly49I (YLI-90), and APC-Ki67 (SolA15), CD45.1 (A20), CD45.2 (104). BUV395-streptavidin was used to develop biotinylated antibodies. Staining of all cells included a preincubation step with unconjugated anti-CD16/32 (clone 2.4G2 or clone 93) mAb to prevent nonspecific binding of mAbs to FcγR. Extracellular staining was performed as described (18). Intracellular staining was performed using the True-Nuclear™ Transcription Factor Buffer Set (BioLegend) per manufacturer’s instructions. Single-color stains were used for setting compensations, and gates were determined by historical data in addition to fluorescent-minus-one control stains. Flow cytometric data were acquired on the BD LSR II flow analyzer or FACS Aria II flow sorter (Becton Dickinson). The experimental data were analyzed using FlowJo™ Software version 10.6.0 and 10.6.3 (Becton, Dickinson and Company).

### Identification of NK cell Ly49 cluster heterogeneity

NK cell clusters (C1-C16) were determined by t-stochastic neighbor embedding (tSNE) using FlowJo™ Software version 10.6.0 and analyzed with 3000 iterations, 50 perplexity, 2800 learning rate (eta), and vantage point tree KNN algorithm (19, 20). The cluster frequencies were determined by manual gating on live, lineage-negative, NK1.1+ cells and further divided by gating on populations based on Ly49I, Ly49G2, Ly49H, and Ly49D expression (please see **Figure 1E**).

**Figure 1.**
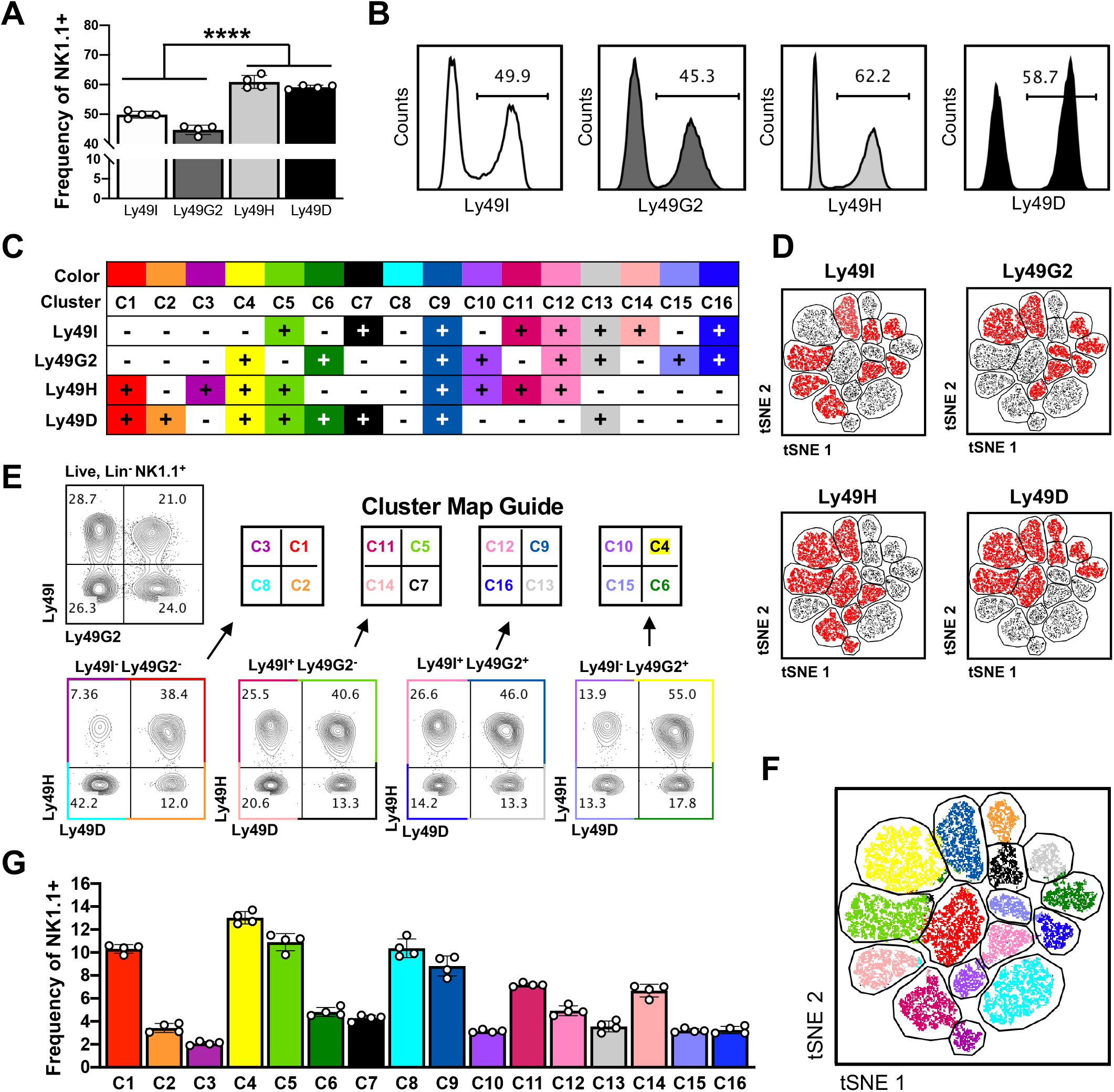
Identification of NK cell cluster phenotypic heterogeneity in B6 mice. (**A**) Frequencies of Ly49I, Ly49G2, Ly49H, and Ly49D receptors on Lin-(negative for CD3, CD4, CD8, CD19, Gr1, and Ter119), NK1.1+ gated cells. Asterisks indicate statistically significant differences between means as determined by the Student’s *t*-test. ****p < 0.0001; (**B**) Representative flow cytometry histogram plots of Ly49 receptor expression shown in A; (**C**) Table of NK cell clusters defined by expression of Ly49 receptors, ordered from most activating (C1) to most inhibitory (C16) based on the number of activating or inhibitory receptors expressed; (**D**) tSNE color mapping plots in red for clusters with Ly49I, Ly49G2, Ly49H, and Ly49D receptor expression; (**E**) Manual gating of 16 NK cell Ly49 clusters gated on live, Lin-NK1.1+ cells; (**F**) Representative unbiased clustering of the 16 clusters visualized by tSNE; (**G**) Frequencies of each cluster in the Lin^−^NK1.1^+^ population. Each point of the graph represents an individual mouse of two separate experiments.

### Assessment of NK cell Ly49 receptor interdependencies by the product rule

Ly49I, Ly49G2, Ly49H, and Ly49D receptors were assessed for independent expression by the product rule. The product rule predicts the frequencies of NK cells that express single or a combination of Ly49 receptors. Assuming a model of independent expression of Ly49 receptors, the frequencies of NK cells expressing individual Ly49 receptors were used to calculate expected frequencies for NK cells expressing zero, one and more receptors (8, 14, 15). For example, the expected frequency for NK cells that co-express Ly49H and Ly49D, but do not express Ly49G2, equals the probability (P) of expressing Ly49H (e.g. frequency of NK cells expressing Ly49H) multiplied by the probability of expressing Ly49D, multiplied by the difference of 1 minus the probability of expressing Ly49G2 (to account for its exclusion), such that P(H,D) = P(H)*P(D)*(1-P(G2)) (please see Figure 2A). We calculated the “product rule error” between the observed and predicted frequencies as the log2(observed frequencies/predicted frequencies). If the observed frequencies were greater than the predicted frequencies (product rule error > 0), this indicated the model underestimated the observed frequencies. Alternatively, if the observed frequencies were less than predicted frequencies (product rule error < 0), this indicated the model overestimated the observed frequencies. If the error equaled zero, then we concluded that the expression of the pattern of receptors were independent. To calculate the total error, we summed the absolute value of the product rule errors amongst clusters.

**Figure 2.**
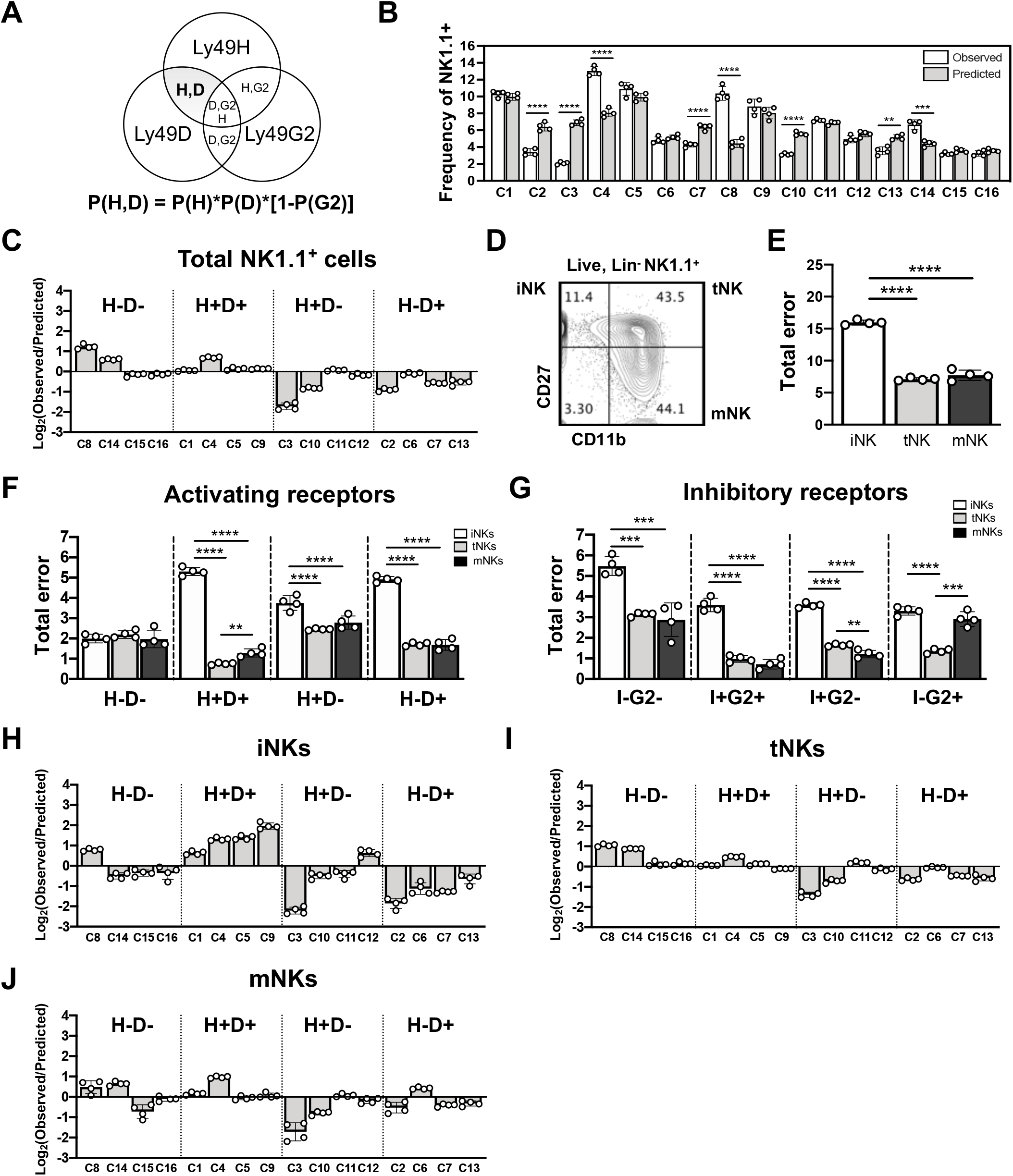
Higher interdependence of NK cell Ly49 cluster frequencies at the iNK stage. (**A**) Example of the product rule calculations of independent probability for NK cells that express Ly49H and Ly49D (H,D) out of three receptors observed. **(B)** Summary graph of experimental (observed) cluster frequencies and cluster frequencies predicted by the product rule (predicted); (**C**) Graph of NK cell log2(observed/predicted) error grouped by activating Ly49 receptors; (**D**) Representative flow cytometry plot of CD27 and CD11b staining to distinguish stages of NK cell maturation; (**E**) Total error was calculated by the sum of the absolute values of log_2_(observed/predicted) from all clusters found at each NK maturation stage determined by the product rule and observed frequencies (iNK, white bars, tNK, gray bars and mNK, black bars); (**F-G**) Total product rule for NK cells organized by expression of (**F**) activating receptors or (**G**) inhibitory receptors; (**H-J**) Log_2_(observed/predicted) error plots for each cluster found in (**H**) iNK, (**I**) tNK, and (**J**) mNK cell stages of maturation. Positive error values indicate underestimation while negative values indicate overestimation. Each point on the graph represents an individual mouse. Asterisks indicate statistically significant differences between means as determined by the Student *t* test. **p < 0.01, ***p < 0.001, ****p < 0.0001.

### In vitro NK cell cluster development assay

Splenic NK cells from C57B6/J mice were harvested and processed to a single-cell suspension in media (RPMI1640 media supplemented with 10% FBS, 0.09 mM nonessential amino acids, 2 mM L-glutamine, 1 mM sodium pyruvate, 100 U/ml penicillin, 100 mg of streptomycin, 0.025 mM beta-mercaptoethanol, and 0.01 M HEPES) and cell counts determined using a hemocytometer. Splenic cells were enriched for NK cells as described in the EasySep Positive Selection Kit (STEMCELL Technologies) manufacturer’s suggested protocol by selecting with an anti-“lineage” cocktail (anti-CD3,-CD4,-CD8,-CD19,-Gr1, and -Ter119). NK cell clusters were then sorted on the FACS Aria II by gating on propidium iodide (PI)-negative (live), “lineage”-negative and NK1.1-positive cells. 90-95% post-sort purity was achieved, as measured by FACS. Approximately 20,000 cells per NK cell cluster were cultured in 96-flat bottom tissue culture plates with the addition of 100,000 lethally-irradiated (30.0 Gy) splenic feeder cells and recombinant-mouse (rm) IL-15 (75 ng/ml). NK cell cluster frequencies were quantified by FACS after 4 days of culture.

### In vivo NK cell cluster adoptive transfer assay

B6.SJL mice were used as a source of donor splenic NK cells. Spleens were aseptically harvested and processed to a single-cell suspension. NK cells were enriched as described above, and then sorted for each of the 16 NK cell clusters. B6 recipient mice were sublethally irradiated (5.5 Gy) with a cesium irradiator four hours before receiving a range of 50,000 to 200,000 donor NK cluster cells by retroorbital intravenous injection. Donor-derived CD45.1+ NK cell cluster frequencies were analyzed by FACS four days after transfer.

### Statistical analysis

Student’s t-test with a two-tailed distribution and with two-sample equal variance (homoscedastic test) was used to determine differences in means between groups using GraphPad Prism software version 8.4.2. A p-value of 0.05 was considered to be statistically significant. Asterisks indicate statistically significant differences *p < 0.05, **p < 0.01, ***p < 0.001, ****p < 0.0001.

## Results

### Identification of NK cell phenotypic heterogeneity amongst the combination of Ly49 receptors

NK cells express diverse frequencies and combinations of Ly49 activating and inhibitory surface receptors, but few studies have examined the relationship between expression of the activating receptor repertoire to that of the inhibitory receptor repertoire, and whether the activating receptors are subject to similar stochasticity in expression as previously described for inhibitory receptors (8, 14). We decided to focus on inhibitory receptors Ly49I and Ly49G2 and activating receptors Ly49H and Ly49D on splenic NK cells in C57B6/J (B6) mice, which express MHC-I ligands H-2K^b^ and H-2D^b^. Ly49I’s inhibitory ligand is H-2K^b^, Ly49H binds to the m157 viral antigen, and Ly49G2 and Ly49D’s known ligand is H-2D^d^ (not expressed in B6 mice) (9). Previous work (10, 17) in which analyses of Ly49H and Ly49D were performed, provided a source of historical controls to which our data could be compared. Although other Ly49 family members exist, we were limited to these 4 receptors due to the availability of fluorochrome-conjugated specific antibodies and the number of fluorochromes available on our flow cytometers. Expression of each Ly49 receptor was observed (**Figure 1A-B**). We defined sixteen NK cell “clusters” (C) by the number and type of activating receptors and inhibitory receptors expressed. That is, Cluster 1 (C1) expresses both Ly49H and Ly49D activating receptors and neither Ly49I nor Ly49G2 inhibitory receptors, whereas Cluster 16 (C16) expresses both Ly49I and Ly49G2 inhibitory receptors but neither Ly49H nor Ly49D activating receptors (**Figure 1C**). By using these four receptors, sixteen possible combinations of Ly49 activating and inhibitory receptor expression were identified by t-distributed stochastic neighbor embedding (t-SNE) analysis of flow cytometric data **(Figure 1F**), and further examination of the expression of individual Ly49 receptors allowed us to confirm each cluster phenotype **(Figure 1D)**(19). Furthermore, we developed an NK cell cluster gating strategy that identified the sixteen unique subpopulations (clusters) and their frequencies **(Figure 1E)**. Similar to previous studies which analyzed inhibitory Ly49 receptors only (8, 14), we observed Cluster 8 (C8), which expresses none of the four receptors, to be one of the most prevalent populations (**Figure 1G)**. By analyzing Ly49 activating receptor frequency, we observed that clusters co-expressing Ly49H and Ly49D activating receptors were most represented, i.e. C1 (I-G2-H+D+), C4 (I-G2+H+D+), C5 (I+G2-H+D+), and C9 (I+G2+H+D+) **(Figure 1G)**. Conversely, frequencies were lowest for clusters that express both Ly49 inhibitory receptors, such as C16 (I+G2+H-D-), C13 (I+G2+H-D+), and C12 (I+G2+H+D-) (**Figure 1G**). These data suggest a possible selective (non-random) pressure controlling the frequencies of these clusters in mice.

### High interdependencies of NK cell cluster frequencies within the immature NK stage

Next, to explore the possibility of independent (random) expression of our Ly49 receptors of interest, we utilized the product rule model for independent expression assuming the stochastic nature of Ly49 receptors. The model states that if Ly49I, Ly49G2, Ly49H, and Ly49D expression is independent of one another, then the observed frequencies of NK cells that express zero, one, two or more receptors can be predicted by the measurement of individual Ly49 receptor frequencies (8, 15) (**Figure 2A**). Thus, if the predicted frequencies match the observed biological frequencies, then we conclude no interdependencies between expression of those Ly49 receptors. Alternatively, if the observed frequencies do not match the predicted frequencies, the expression of these receptors are interdependent (non-random) for reasons including, but not limited to, linked receptor expression, gene regulation, and biased receptor selection (8, 10, 21). Our comparison of the observed versus predicted NK cluster frequencies demonstrate independent expression in some clusters (C1, C5, C6, C9, C11, C12, C15 and C16), but dependencies in others (C2, C3, C4, C7, C8, C10, C13 and C14) (**Figure 2B**). We next calculated the product rule error (e.g. log2(observed/predicted)) for each cluster to identify if the model underestimated (observed > predicted), overestimated (observed < predicted), or matched the observed frequencies (observed = predicted). An example of the product rule errors in clusters arranged by their patterns of Ly49H and Ly49D expression is shown in **Figure 2C**. We show that NK clusters C14, C8 (which are H-D-), and C4 (H+D+) frequencies were underestimated by the model, whereas clusters C10, C3 (both H+D-) and, C7, C13, and C2 (H-D+), were overestimated by the model (**Figure 2C**). We noted that the product rule underestimated the frequencies of five out of the eight clusters that express only one of the Ly49 activating receptors (C10, C3, C7, C13, and C2). These data suggest interdependencies between Ly49H and Ly49D expression.

We hypothesized that a source of NK cell Ly49 expression dependencies was the stage of maturation. To test our hypothesis, we utilized the NK cell maturation markers CD27 and CD11b to examine the frequencies of each cluster amongst immature NK (iNK; CD27^+^CD11b^−^), transitional NK (tNK; CD27^+^CD11b^+^), and mature NK (mNK; CD27^−^CD11b^+^) cells (1, 18, 22, 23) (**Figure 2D)**. We calculated the total error by summing the absolute values of the product rule errors for each NK cell maturation stage (**Figures 2E, 2H-J**). The total error was significantly higher at the iNK cell stage relative to the tNK and mNK cell stages, suggesting more Ly49 receptor dependencies at the iNK stage (**Figure 2E**). iNK total error was significantly higher in clusters which at least one activating receptor (**Figure 2F**). Examination of the inhibitory receptors showed very similar patterns (**Figure 2G**). The exceptions were H-D-clusters which displayed similar error between iNK, tNK, and mNKs, and I-G2+ clusters, in which error was high in the iNK and mNK cells (**Figure 2F-G**). Moreover, C8, C14, C15, and C16, all of which are H-D-, showed more independent expression (lower total error) in the iNK (1.99 ± 0.21) and tNK (2.20 ± 0.18) cells relative to I-G2-clusters C1, C2, C3, C8 in the iNK (5.48 ± 0.45) and tNK (3.1 ± 0.10) cells (**Figure 2F-G**). No difference between the total error for tNK and mNK cells within the H-D- and I-G2-groups were observed (**Figure 2F-G**). Additionally, we observed the majority of clusters were either overestimated or underestimated consistently throughout NK cell maturation (**Figure 2H-J**). There were some exceptions: C14 (I+G2-H-D-) was overestimated by the model at the iNK stage and then underestimated at the tNK and mNK maturation stage, and C6 (I-G2+H-D+) was overestimated at the iNK stage and then slightly underestimated at the mNK cell stage (**Figure 2H-J**). Taken together, the product rule model provided evidence for a tightly regulated expression system for Ly49 receptors especially at the iNK maturation stage, with the highest interdependencies in clusters that express at least one Ly49H and Ly49D activating receptor.

### Dominant expression of Ly49I in all clusters at the mature NK cell stage

Further analysis revealed Ly49 cluster frequencies were unique at each NK cell maturation stage (**Figure 3A-D**). We identified Ly49I receptor expression frequency to significantly increase from iNK to tNK to mNK stages (**Figure 3E**). Ly49G2, Ly49H, and Ly49D frequencies significantly increased between iNK to tNK stages, and then decreased (Ly49G2 and Ly49D) or stabilized (Ly49H) moving from the tNK to mNK stages (**Figure 3E**). The frequencies of clusters with zero or one receptor were found to decrease; conversely, clusters expressing two, three, or four receptors increased from iNK to tNK stages (**Figure 3F**).

**Figure 3.**
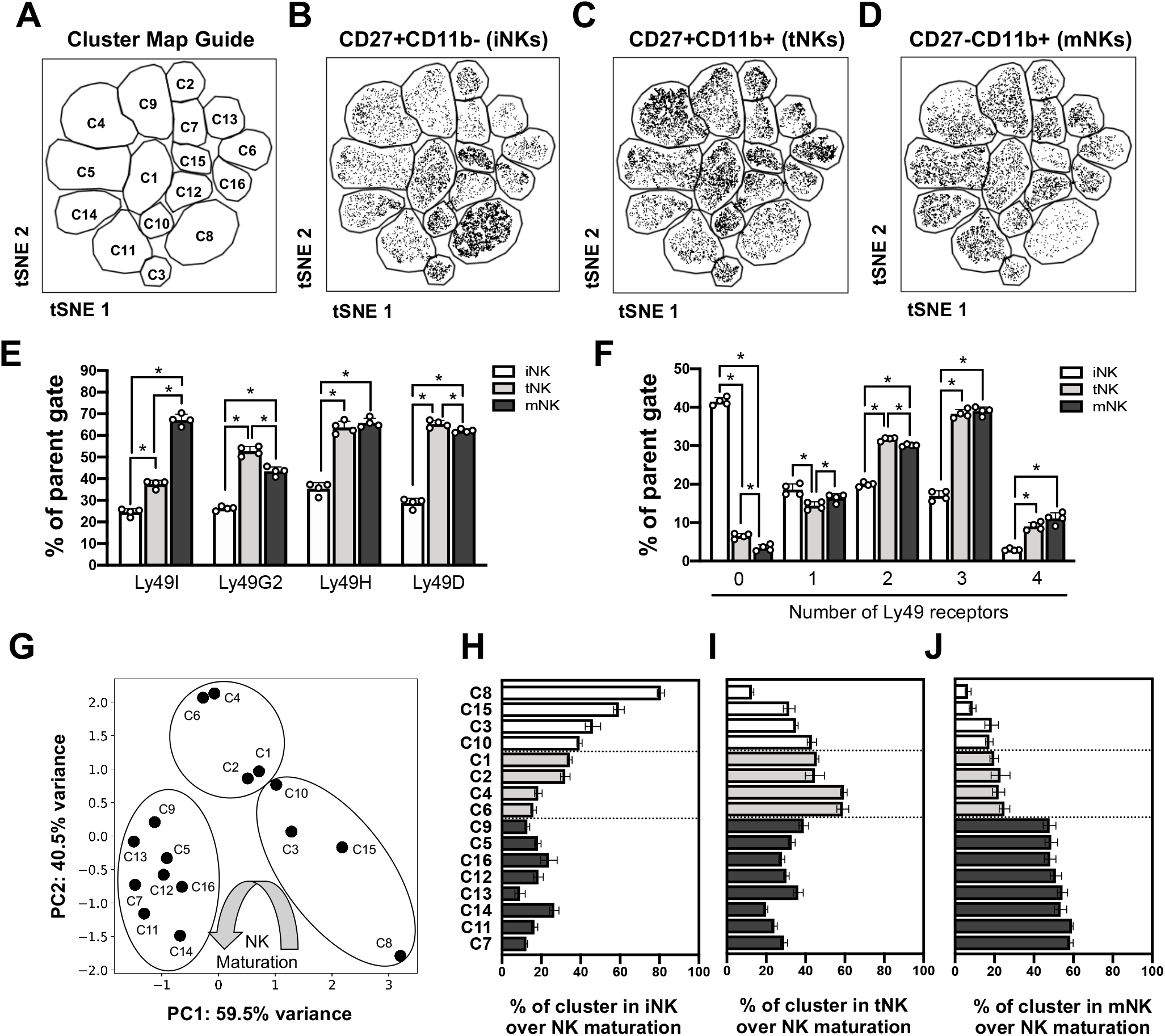
Certain NK clusters predominate at each stage of NK cell maturation. (**A**) Map of cluster location via tSNE plot; (**B-D**) Representative tSNE plots of (**B**) iNK, (**C**) tNK, and (**D**) mNK cells visualized by density; (**E**) Summary graph of Ly49I, Ly49G2, Ly49H, and Ly49D receptor frequencies gated on iNK (white bars), tNK (gray bars), or mNK cells (black bars); (**F**) Summary graph of each NK cell maturation stage expressing 0, 1, 2, 3, or 4 Ly49 receptors; (**G**) PCA plots were generated from 4 biological replicates of average percentage for clusters with respect to NK cell maturation stage; (**H-J**) Summary graph of each calculated cluster percentage for each cluster at (**H**) iNK, (**I**) tNK, (**J**) and mNK cell stage. Each point of the graph represents an individual mouse. Asterisks indicate statistically significant differences between means as determined by the Student *t* test. *p < 0.05.

To determine if specific clusters were predominantly grouped at the iNK, tNK or mNK cell stages of maturation, we quantified the cluster frequencies at each NK maturation stage and visualized the similarities between clusters with principal component analysis (PCA) (24). NK cell cluster percentages were quantified by normalizing each cluster frequency relative to each NK maturation stage, such that a cluster’s frequency was divided by the sum of that cluster’s frequencies found at each maturation stage and then multiplied by 100 (**Supplemental Figure 1**). We computed the PCA from the average normalized frequencies of clusters at each NK stage and determined similarities between clusters with respect to NK maturation (**Figure 3G-J**). We grouped clusters found predominantly at the iNK, tNK, and mNK cell stage in our PCA, which were confirmed by observing the plotted percentages (**Figure 3H-J**). That is, we identified C8, C15, C3, and C10 to predominate the iNK cell stage (**Figure 3H**), C1, C2, C4, C6 to predominate at the tNK cell stage (**Figure 3I**), and C9, C5, C13, C16, C7, C11, C14 to predominate at the mNK cell stage (**Figure 3J**). C10, C1 and C2 assembled close together in the PCA, but we decided to group C10 into the “iNK-predominant” group because the changes in the % of C10 at the iNK, tNK and mNK were more similar to to C8, C3 and C15 (**Supplemental Figure 1**). Notably, the predominant clusters within each stage were represented at different proportions as maturation progressed. For example, C8, which expresses none of the four receptors, decreased in a sequential manner throughout NK cell maturation from 80.7% ± 1.8 in the iNK stage to 6.6% ± 1.5 at the mNK cell stage (**Figures 3H and 3J**). In contrast, tNK-predominant clusters were lower at the iNK and mNK stages, and mNK predominant clusters were lowest at the iNK and tNK stages (**Figures 3H, 3I, 3J).** Furthermore, all mNK-predominant clusters expressed the inhibitory self-Ly49I receptor, but iNK- and tNK-predominant clusters did not (**Figures 1C, 3H, 3I and 3J**) Thus, these findings suggest that the frequency and phenotype of the NK cell clusters are regulated throughout NK cell maturation to increase the types of receptors expressed, and positively select for self-Ly49I at the mNK cell stage.

### Evidence for prescribed NK cell Ly49 developmental pathways in mice

The observed distribution of clusters within the iNK, tNK and mNK stages (**Figure 3 and Supplemental Figure 1**) led us to hypothesize that the iNK-predominant clusters may be precursors to clusters that predominate at the tNK to mNK stages. To test this, we sorted each of the 16 NK cell clusters and cultured them with 75 ng/ml recombinant-mouse-interleukin-15 (rmIL-15) and lethally irradiated splenic feeder cells for 4 days (**Figure 4A**). We observed that iNK-predominant clusters C8, C15, C3, C10, and tNK-predominant clusters C1, C2, C4, C6, all upregulated Ly49I receptor after culture (**Figure 4B, 4C and 4D**), which would re-categorize them into clusters found predominantly in the mNK cell stage (C5, C7, C9 and C13, respectively, **Figures 1C and 3J**). Cultures initiated with mNK-predominant clusters maintained their Ly49 receptor phenotypes (data not shown). To verify our findings in vivo, we adoptively transferred sorted C8, C15, C3, C10, C1, C2, and C14 cells from B6.SJL (CD45.1+) mice into sublethally irradiated B6 (CD45.2+) hosts and analyzed their differentiation after 4 days (**Figure 4E**). Similar to our in vitro results, we observed C8, C15, C3, C10, C1, and C2 to upregulate Ly49I **(Figure 4F-I**), and that Ly49H and Ly49D maintained stable expression (data not shown). However, in vivo, C8, C1, C2 upregulated a small frequency of inhibitory Ly49G2 receptor after 4 days (**Figure 4F and 4G**). Ly49 receptor expression on C14, a mNK-predominant cluster, was unchanged after adoptive transfer (**Figure 4H).** Altogether, these results strongly support the existence of prescribed pathways of NK cell maturation from precursor NK cell clusters, and that Ly49I upregulation is a key step for mature NK cells (**Figure 4J and 4K**).

**Figure 4.**
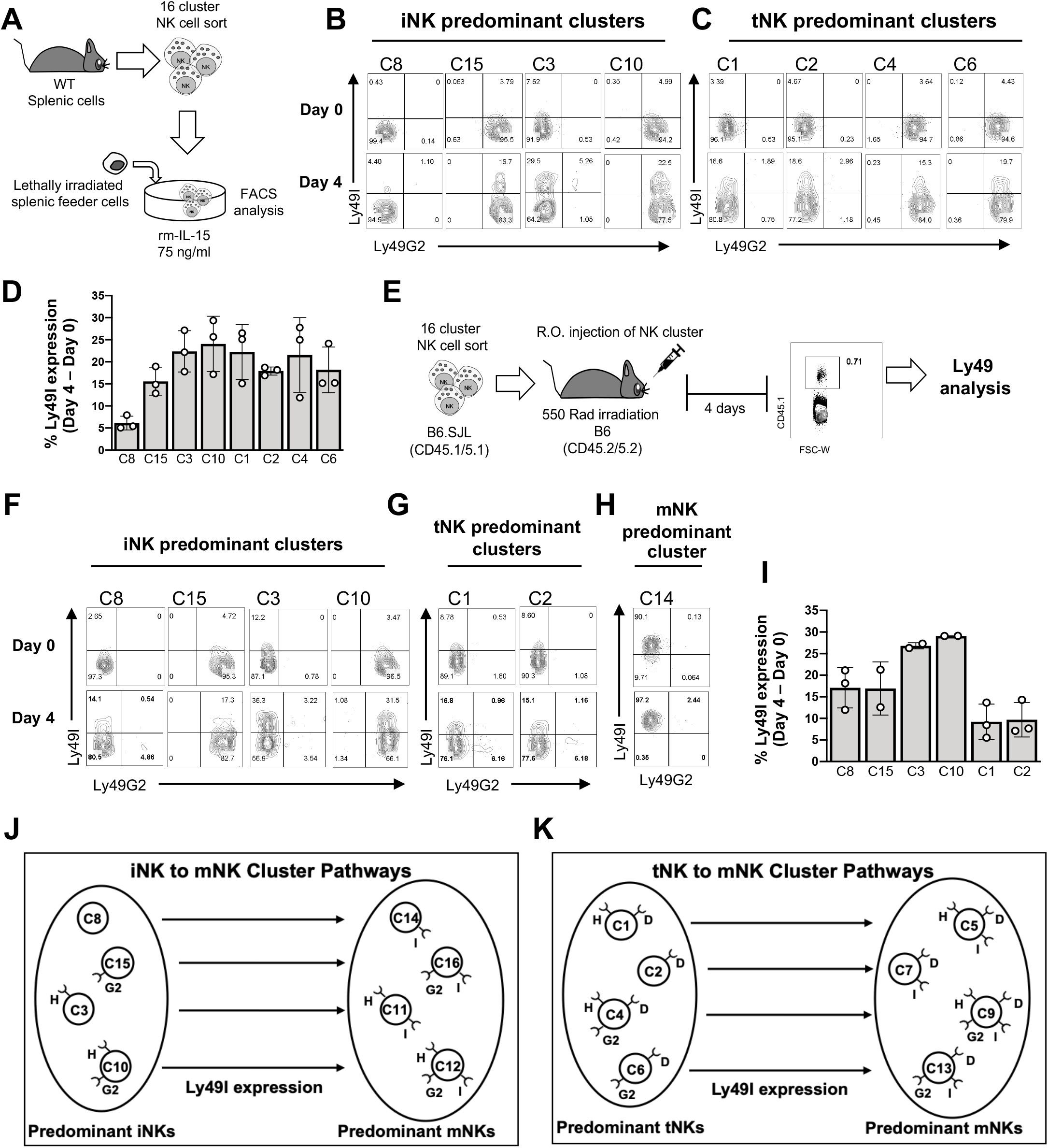
Evidence for prescribed pathways of NK cell cluster differentiation in vitro and in vivo. (**A**) Scheme of experimental design for sorting clusters and in vitro culture; (**B-C**) Representative Ly49I vs. Ly49G2 flow cytometry plots of sorted clusters showing post-sort purity on Day 0 (top row) and 4-days after culture (bottom row) of gated NK1.1+ cells; (**B**) Differentiation of clusters predominantly found at the iNK stage, (**C**) differentiation of clusters predominantly found at the tNK stages: (**D**) Summary graph of cluster Ly49I receptor expression after Day 4 of culture, calculated by subtracting Day 4 Ly49I frequency from Day 0 Ly49I frequency; (**E**) Scheme of experimental design of in vivo adoptive transfer of selected NK clusters; (**F-H**) Representative Ly49I vs. Ly49G2 flow cytometry plots of sorted clusters for post-sort purity of gated NK cells on Day 0 (top row) and Day 4 after adoptive transfer into recipient mice (bottom row); (**F**) In vivo differentiation of clusters predominantly found at the iNK stage; (**G**) In vivo differentiation of clusters predominantly found at the tNK stage, and (**H**) in vivo differentiation of C14, found predominantly in mNK; (**I**) Summary graph of cluster Ly49I receptor expression 4 days after adoptive transfer, calculated as in D. Each point of the graph represents an experimental replicate; (**J**) Schematic summary of prescribed pathways from iNK-predominant clusters; (**K**) Summary of prescribed pathways from tNK-predominant clusters.

### Immature NK cells display similar proliferation characteristics, regardless of cluster type

Given our discovery of prescribed pathways of NK cluster development, we further analyzed the clusters for differences in their proliferative state. We hypothesized that the proliferation rates of specific NK cell clusters would be distinct. Furthermore, we expected proliferation rates to correlate with maturation stage. To test our hypotheses, we sorted and cultured bulk iNKs, tNKs, and mNKs (**Figure 5A**) on lethally irradiated splenic feeder cells in media containing 75 ng/ml of rm-IL-15, and measured cellularity at Day 2 and Day 6 post-culture. We observed the highest fold change in cellularity for cultures initiated with iNK cells, as compared to cultures initiated with tNK or mNK cells (**Figure 5B**). We disaggregated the fold changes in proliferation between iNK-predominant (C8, C15, C3, C10), tNK-predominant (C1, C2, C4, C6), and mNK-predominant (C9, C5, C16, C12, C13, C14, C11, C7) clusters within each sorted population **(Figure 5C, 5D, 5E)**. Our data show that iNK-initiated cultures increased the fold change in cellularity of tNK-predominant clusters relative to iNK- and mNK-predominant clusters (**Figure 5C**). In contrast, mNK-initiated cultures showed reduced proliferation in tNK predominant clusters relative to iNK- and mNK-predominant clusters (**Figure 5E**). Additionally, the frequency of C8, which expresses none of the four Ly49 receptors and is the most prevalent in the iNK stage, dramatically decreased during the culture period **(Figure 5F)**, which is consistent with our observation of decreased frequency of C8 at the tNK and mNK stages in vivo **(Figure 3I and 3J**).

**Figure 5.**
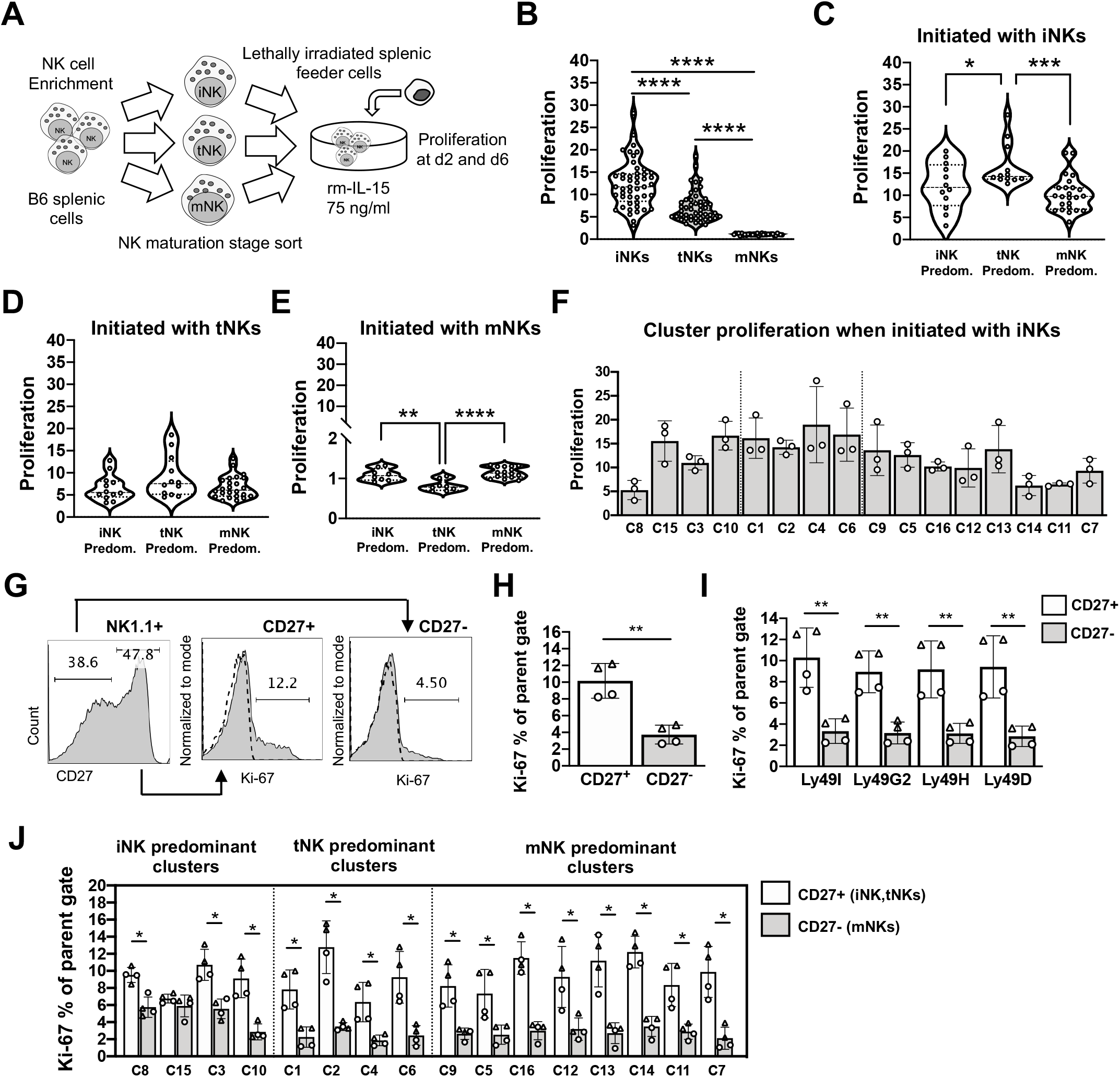
Immature NK cells display in vitro characteristics of proliferation, regardless of cluster type. **(A**) Scheme of experimental design to sort and culture iNK (CD27^+^CD11b^−^), tNK (CD27^+^CD11b^+^), and mNK (CD27^−^CD11b^+^) cells; (**B-E**) NK cell proliferation as determined by flow cytometry and calculating the fold change in cellularity between Day 2 and Day 6 of culture (e.g. # of cells at Day 6 divided by # of cells at Day 2) for (**B**) overall cell proliferation, proliferation in cultures initiated with sorted (**C**) iNKs, (**D**) tNKs, and (**E**) mNKs, further categorized into predominant iNK, tNK, and mNK clusters (n=3 experimental replicates; (**F**) Fold change in proliferation for clusters that originated from sorted iNK cells; (**G**) Intracellular Ki-67 expression of Lin^−^NK1.1^+^ CD27^+^ (iNK and tNK stages) and Lin^−^NK1.1^+^ CD27^−^(mNK stage) cells post-culture (n=2 female mice (triangles; 24 weeks old), n=2 male mice (circles; 19 weeks old); (**H**) Summary plots of Ki-67 expression between CD27^+^ or CD27^−^NK cells: (**I**) Ki-67 expression on CD27^+^ and CD27^−^NK cells disaggregated by Ly49 receptor type, and (**J**) Ki-67 expression on NK cells disaggregated by cluster type. Asterisks indicate statistically significant differences between means as determined by the Student*’s t* test. *p < 0.05, **p < 0.01, ****p < 0.0001.

To confirm these proliferation patterns, we stained NK cells for Ki-67 expression post-culture. After culture, we noticed that CD11b expression was downregulated universally on NK cells, preventing us from using CD11b to distinguish iNK, tNK and mNKs (data not shown), likely as an artifact of in vitro culture (25). However, CD27 expression was still binary, allowing us to distinguish CD27+ (presumably iNK and tNKs) from CD27-mNKs. We found that NK1.1^+^CD27^+^ cells expressed higher Ki-67 levels compared to NK1.1^+^CD27^−^cells (**Figure 5G and 5H**), regardless of Ly49 receptor expression (**Figure 5I**). This pattern persisted when the Ki-67 expression was examined in specific NK cell clusters, with the exception of cluster C15 (which only expresses Ly49G2) (**Figure 5J**). These data show that cluster designation (and hence Ly49 receptor expression) does not dictate NK cell proliferation. Rather, proliferative potential is a general characteristic of NK cell maturation stage, with highest proliferation in the iNK cells and lowest proliferation in mNK cells.

### MHC-I-deficiency does not affect NK cell maturation, but results in underrepresentation of NK cell clusters which express activating Ly49 receptors

We next investigated the effects of MHC-I on NK cell cluster heterogeneity and NK cell maturation in β2m^−/−^(MHC-I^−/−^) mice. In MHC-I^−/−^spleens, the frequencies of NK cells expressing inhibitory Ly49I and Ly49G2 receptors increased, whereas the frequencies of NK cells expressing activating Ly49H and Ly49D receptors decreased (**Figure 6A**). These differences in Ly49 frequencies in MHC-I^−/−^mice did not appear to influence NK cell maturation (**Figure 6B**). MHC-I^−/−^mice displayed significantly lower frequencies of I-G2-NK cells (**Figure 6C and 6D**) and higher frequencies of H-D-NK cells (**Figure 6E and 6F**). Thus, NK cells from MHC-I^−/−^mice have decreased frequencies of activating receptors and increased frequencies of inhibitory receptors. The increased frequency of inhibitory receptor expressing NK cells is due to an increase in dual inhibitory receptor expressing I^+^G2^+^ NK cells, as lower frequencies of I-G2+ NK cells were observed in MHC-I^−/−^mice (**Figure 6C-D**). Additionally, our activating receptor analysis showed that H+D-NK cells increased, whereas H+D+ cell frequencies decreased in MHC-I^−/−^mice (**Figure 6E-F**). Further investigation of individual Ly49 receptor frequencies as a function of iNK, tNK and mNK maturation stages revealed different behaviors of the inhibitory and activating Ly49 receptor behaviors in MHC-I^−/−^mice (**Figure 6G and 6H**). The frequencies of Ly49I+ and Ly49G2+ NK cells were significantly higher in MHC-I^−/−^mice compared to MHC-I^+/+^ (wild-type) controls at each stage of NK maturation (**Figure 6G**). In contrast, the frequencies of Ly49H+ and Ly49D+ NK cells were significantly decreased only at the tNK and mNK stages, but similar at the iNK stage (**Figure 6H**). Overall, these data suggest MHC-I molecules play a differential role in the expression of inhibitory and activating Ly49 receptors, and that MHC-I is not essential for progression from iNK, tNK, and mNK stages during maturation.

**Figure 6.**
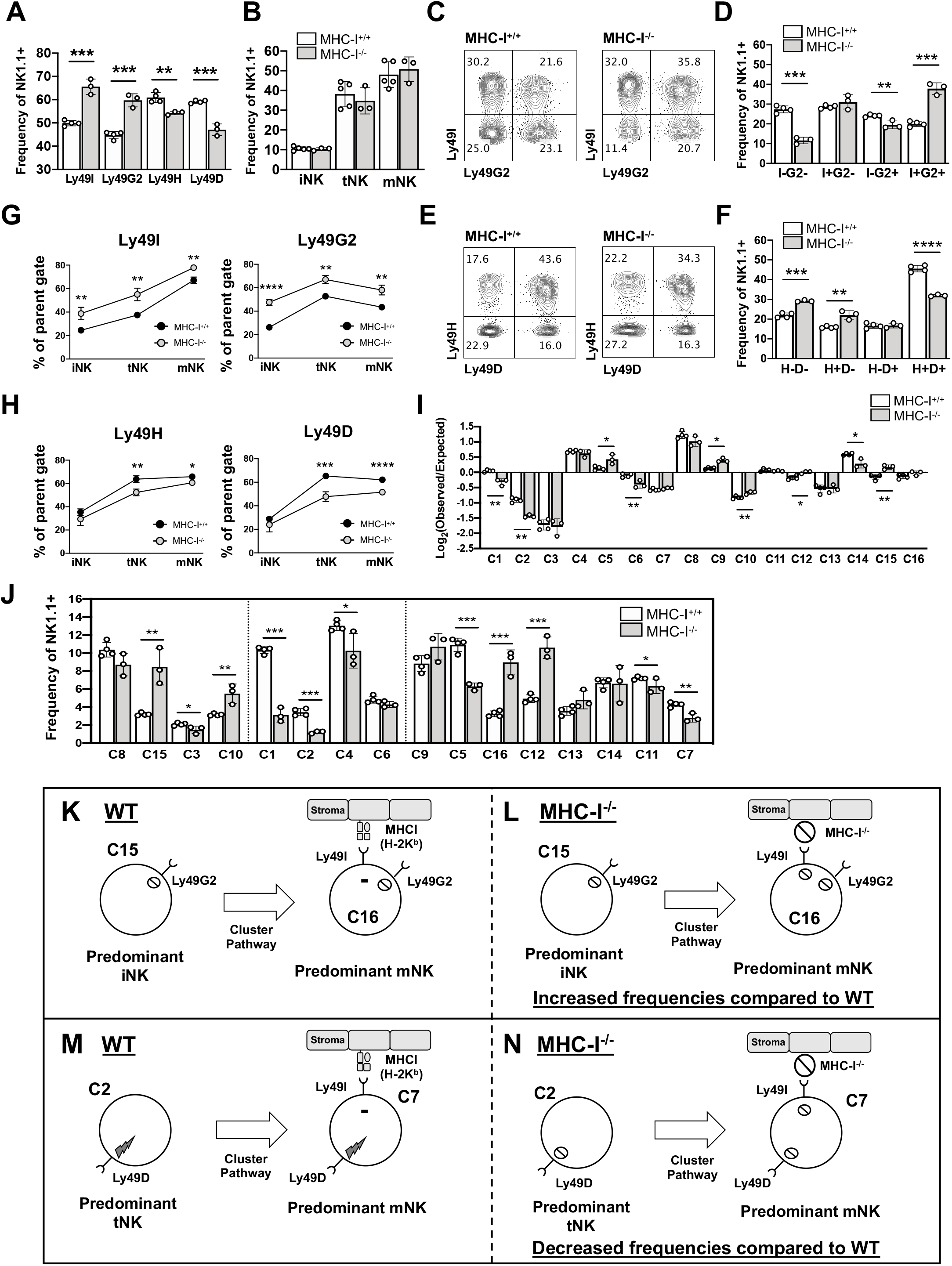
Altered NK cell cluster distribution in MHC-I^−/−^mice. (**A**) Summary graph of Ly49I, Ly49G2, Ly49H, and Ly49D frequencies gated on NK1.1+ cells in β2m^−/−^(MHC-I^−/−^) and B6 (MHC-I^+/+^) spleens; (**B**) Graph of frequencies of iNK (CD27^+^CD11b^−^), tNK (CD27^+^CD11b^+^), and mNK (CD27^−^CD11b^+^) cells; (**C,E**) Representative plots of Lin^−^NK1.1^+^ cells examining (**C**) Ly49I and Ly49G2 and (**E**) Ly49H and Ly49D in MHC-I^+/+^ (left) and MHC-I^−/−^mice; (**D,F**) Summary plots for data shown C and E; (**G**) Graphs gated on parent iNK, tNK or mNK gate for Ly49I and Ly49G2; (**H**) Graph gated on parent iNK, tNK or mNK gate for Ly49H and Ly49D; (**I**) Log2(observed/expected) values for MHC-I^−/−^and MHC-I^+/+^ NK cell clusters; (**J**) Graph of NK cell cluster frequencies in MHC-I^+/+^ and MHC-I^−/−^mice; (**K-N**) Working model illustrating MHC-I’s influence on NK cell pathways for clusters expressing only inhibitory receptors (**K-L)** or activating receptors (**M-N**); (**K and M**) model of cluster pathways in MHC-I^+/+^ mice; (**L and N**) model of cluster pathways in MHC-I^−/−^mice. Each point of the graph represents an individual mouse. Asterisks indicate statistically significant differences between means as determined by the Student *t* test. *p < 0.05, **p < 0.01, ***p < 0.001, ****p < 0.0001.

We next compared NK cell cluster Ly49 receptor expression dependencies amongst WT and MHC-I^−/−^mice using the product rule. Any deviations between the two conditions (WT and MHC-I^−/−^) would indicate dependencies which are influenced by MHC-I expression. We compared the product rule errors for each cluster in WT and MHC-I^−/−^mice, and observed significantly different errors in nine of the sixteen clusters in MHC-I^−/−^mice (**Figure 6I**). C1, C2, C5, C6, and C9 were found to increased error (e.g. more dependencies) in MHC-I^−/−^mice, and these clusters commonly express the activating Ly49D receptor (**Figure 6I**). In contrast, C10 (I-G2+H+D−), C12 (I+G2+H+D−), and C14 (I+G2−H−D−) in MHC-I^−/−^mice decreased dependencies (error) relative to WT, and do not express Ly49D. Furthermore, C15 (I−G2+H−D−) was the only cluster that flipped directionality of the error from being overestimated in WT to underestimated in MHC-I^−/−^mice. This suggest that Ly49G2 is negatively regulated in MHC-I sufficient microenvironments.

Next, we wanted to determine MHC-I’s influence on NK cell cluster distributions. First, we examined NK cell frequencies based on the number of Ly49 receptors expressed, regardless of receptor type. Collectively, clusters that expressed only one receptor were increased at the iNK cell stage (**Supplemental Figure 2A**), but this increase was attributed to an increase in C15 only (**Figure 6J**). The frequencies of clusters which express two receptors were collectively lower at all stages of NK cell maturation in MHC-I^−/−^mice (**Supplemental Figure 2B)**, which was attributed to decreases in C1, C6, C7, and C11 clusters. C10 and C16, which also expressed 2 receptors, were increased in MHC-I^−/−^mice (**Figure 6J**). The overall increase in NK cells that express 3 receptors in MHC-I^−/−^mice was attributed to C12 (**Supplemental Figure 2C, Figure 6J**). All of the clusters that were increased (C15, C10, C16, and C12) express Ly49G2 (**Figure 6J**). Furthermore, C3, C1, C2, C5, C7, and C11, which lack Ly49G2 expression, were significantly decreased in NK cell frequencies (**Figure 6J**). These data suggest that MHC-I is a major regulator of Ly49G2 expression, despite the fact that no known MHC-I ligand for Ly49G2 in B6 mice has been described (3). In addition, Ly49G2-positive clusters C15, C10, C16, and C12 all lack Ly49D expression and were increased. In contrast, the Ly49G2-negative clusters C1, C2, C5, and C7, which express Ly49D, were decreased (**Figure 6J**). This suggests that there is a reciprocal dependency between Ly49D and Ly49G2 expression and MHC-I in the distribution of cluster frequencies. No common relationship between MHC-I deficiency and Ly49I or Ly49H on cluster distributions could be determined with our cluster frequency data set (**Figure 6J**).

We continued to investigate the influence of MHC-I on NK cell cluster frequencies found predominantly within the iNK, tNK, and mNK cell stage of maturation. Similar to the previous analysis in which we determined the predominant clusters within NK maturation stages (**Figure 3**), we compared MHC-I^−/−^and WT (MHC-I^+/+^) clusters using PCA. We observed that most clusters in MHC-I^−/−^matched the same trends in cluster maturation found in WT mice (**Supplemental Figure 2D)**; however, C1, C2, and C16 maintained steady frequencies throughout maturation in MHC-I^−/−^mice (**Supplemental Figure 2F and 2G**). In WT mice, C15, C10, and C3 iNK-predominant clusters transition into C16, C12, and C11 (**Figure 4**). In MHC-I^−/−^mice, we observed increased C15 and C10 frequencies, matching the observed frequencies for C16 and C12. Furthermore, we observed decreased C3 and C11 frequencies (**Supplemental Figure 2E and 2G**). Additionally, tNK-predominant clusters C2 and C1 were both decreased in MHC-I^−/−^mice and their subsequent mNK clusters C7 and C5 were also both decreased (**Figure 6J**). Although tNK C4 was decreased, its subsequent cluster, C9, was found in normal frequencies at the mNK stage. C6 to C13 frequencies were also unaffected. Notably, C4, C9, C6 and C13 all co-express Ly49G2 and Ly49D. Overall, these data lend further support to the prescribed pathways of NK cluster differentiation and suggest that MHC-I influences the cluster frequencies via regulation of Ly49G2 and Ly49D expression.

## Discussion

Previously, the product rule has been used to determine the interdependencies of inhibitory Ly49 receptors and their respective MHC-I ligand (8, 15). In this study, we extended this analysis to include the Ly49I inhibitory receptor in combination with activating receptors Ly49H and Ly49D, as well as a careful study of Ly49 receptor “clusters” based on NK cell maturation. Here, we show for the first time that the majority of the interdependencies in Ly49 receptor expression originate at the iNK cell maturation stage (8, 14), and that specific clusters predominate at each stage. Our results demonstrate strong interdependencies of the activating Ly49H and Ly49D receptors, which we propose can explain the altered cluster frequencies observed in MHC-I^−/−^mice. Moreover, our results further resolve a role for Ly49I as an important selective marker of completion of NK cell development in B6 mice (22).

Ly49I’s known ligand in B6 mice is MHC-I K^b^ (3, 9, 26). Our NK cluster development studies suggest a process in which Ly49I-negative iNK and tNK clusters develop with high proliferative potential. We propose that when Ly49I is upregulated, transition into the mNK cell stage occurs, and proliferation is then inhibited by interactions between Ly49I and K^b^. These observations are most consistent with the “sequential expression model”, which proposes splenic NK cells sequentially accumulate inhibitory Ly49 receptors until receptors specific for self MHC-I molecules are expressed (14, 27, 28). We observed that Ly49I+ clusters become more prevalent overall at the mNK cell stage, indicating that NK cells also sequentially accumulate activating and non-self-binding Ly49 receptors before expressing Ly49I. Consistent with previous findings, we observed decreased mNK cell proliferation relative to iNK and tNK cells (**2**), which may suggest that self-inhibitory Ly49I completes the maturation process and maintains mNK cells in a quiescent state ready to be triggered in an immune response.

Our data expand the sequential expression model and suggest an updated working model in which MHC-I affects NK cell Ly49 activating and inhibitory receptor expression and alters the prescribed pathways of NK cluster differentiation, but via distinct mechanisms. Our model distinguishes between the differentiation of clusters expressing only inhibitory (**Figure 6L-M**), or only activating receptors at an early maturation stage (**Figure 6N-6O**). In this working model, we focus on the Ly49 receptor interactions at the iNK and tNK stages of maturation that result in upregulation of Ly49I to complete NK cell maturation. In **Figure 6L**, in the predominant iNK cluster that expresses inhibitory Ly49G2 but no activating receptors (C15), Ly49G2 inhibitory signals are not initiated (because there is no Ly49G2 ligand in B6 mice). Due to this lack of inhibitory signal, developing NK cells then upregulate inhibitory receptor Ly49I (differentiating to C16). Ly49I binds to its H-2K^b^ ligand, and in turn, C16 NK cell development is completed and then sustained (**Figure 6L**). The increased frequencies of C15 and C16 observed in MHC-I^−/−^mice (**Figure 6I**) can be explained by this model (**Figure 6M**). In MHC-I^−/−^mice, the C15 cluster upregulates Ly49I in the same fashion as in WT mice. However, Ly49I inhibitory signals are not generated (because H-2K^b^ is not present) and the developing C15 cluster continues to expand (**Figure 6M**). Extending this working model to clusters that co-express a single activating Ly49 receptor with Ly49G2 suggests that Ly49H and Ly49D regulate cluster frequencies differently, as C10 (G2+H+D-I-) and its mature counterpart C12 (G2+H+D-I+) are both increased in MHC-I^−/−^mice, whereas frequencies of C6 (G2+D+H-I-) and its counterpart C13 (G2+D+H-I+) are unaffected. This suggests that the Ly49H+ clusters are controlled by presence of MHC-I. However, C4, which expresses both activating receptors (G2+H+D+I-), was lower in frequency in MHC-I^−/−^mice, but frequencies of its counterpart C9 (G2+H+D+I+) were normal. This indicates that the roles of Ly49H and Ly49D during NK cell development are complex, and MHC-I is not essential for some of these roles.

Our accompanying working model of NK cell development starting with clusters expressing only activating receptors at the iNK and tNK stages is shown in **Figure 6N**. The known ligand for Ly49H is m157, a mouse cytomegalovirus MHC-like protein (2, 29), while the known ligand for Ly49D is H-2D^d^. No known self-MHC-I ligand has been identified for Ly49H and Ly49D in B6 mice (30), but it is possible weak binding to self-MHC-I or non-MHC-I ligands exist (31, 32). Freund et al. reported that activation signals via SLP-76 upregulates inhibitory Ly49A, Ly49G2, and Ly49I receptor expression (10). Similarly, we presume that in B6 mice, an activating ligand exists for Ly49H and Ly49D. NK clusters with one activating receptor (e.g. C2, C3) will generate a signal in response to this activating ligand, which promotes its differentiation and expansion. The activating signal also results in upregulation of inhibitory Ly49I at the mNK stage (**Figure 6N**). We interpret the clear effects of MHC-I deficiency on Ly49H and Ly49D frequencies to demonstrate a relationship between MHC-I and the activating receptors, but that this is not a direct ligand-receptor interaction. NK cell development and NK cell survival in MHC-I-deficient mice may be impaired by dysfunctional dendritic cell expression of IL-12 and IL-15 transpresentation (33, 34). In MHC-I^−/−^mice, we assume the activating ligand is absent or signaling is impaired. In the absence of these signals, low expansion and impaired differentiation to the corresponding mNK cluster in MHC-I^−/−^mice results. Lack of activating signals in MHC-I^−/−^also dysregulates the expression of Ly49I at the mNK stage (**Figure 6O**). We observed an enhanced decrease in cluster frequencies in MHC-I^−/−^mice when two activating receptors were expressed (C1 and C5), which we posit could result from lower expansion of dual H+D+ clusters (**Figure 6I**). However, C9, which is H+D+ and also co-expresses both inhibitory receptors, is unaffected in the MHC-I^−/−^, suggesting a “canceling out” or “balancing” of activating and inhibitory signals in this case. This balancing is further supported by the similar frequencies of C6 and its mNK counterpart C13, which express one inhibitory and activating receptor, in B6 and MHC-I^−/−^mice (**Figure 6I)**.

In our study, we focused on the role of Ly49 receptors on NK cell development. However, given that NK cytotoxicity is governed by balance of signals between Ly49 activating and inhibitory receptors **(3)**, it is also possible that our evidence of prescribed NK cell developmental pathways can be applied to the identification of NK cell clusters with high cytotoxic potential. Transcription of cytotoxicity genes increases with NK maturation with the highest cytotoxic gene expression at the mNK stage (22, 23). NK cell licensing via the expression of Ly49I and its binding to K^b^ is consistent with our observations that mNK cells acquire Ly49I expression in a manner that correlates to NK cell maturation.

Although we have focused on splenic NK cells in our work, it is also noteworthy that mice and human NK cell maturation are could be tissue-specific, which may change the NK cell Ly49 cluster pathways we observed in the spleen (22, 35). Although NK cells originate in the bone marrow, they continue to mature in peripheral tissues. However, these tissues have been shown to express different frequencies of iNK, tNK, and mNK cells (22, 36). The bone marrow and lymph node tend to express high frequencies of iNK and tNK cells, whereas the lung compartment expresses mostly mNK cells. The spleen, liver and blood tend to express all three stages of NK cell maturation (22). Additional studies are necessary to determine if the prescribed pathways of NK development we observed in the spleen are conserved in other tissues.

## Acknowledgments

We thank the staff of the Department of Animal Research Services and the Flow Cytometry Core of the Stem Cell Instrumentation Foundry at the University of California Merced for excellent animal care and technical support. We thank Dr. Kirk Jensen with the gift of rm-IL-15 and Dr. Anna Beaudin for sharing flow cytometry reagents. We are also grateful to Drs. Marcos E. García-Ojeda and Katrina K. Hoyer, as well as the UC Merced Immunology Journal Club for their comments on the manuscript.

## Abbreviations

b2m: beta-2-microglobulin
C: cluster
iNK: immature NK
Lin: Lineage
MHC-I: MHC class I
mNK: most mature NK
PCA: principal component analysis
tNK: transitional NK
tSNE: t-stochastic neighbor embedding

