## Supplemental Figures for "Evidence for prescribed NK cell Ly49 developmental pathways in mice"

### Millan et al. 2020. Supplemental Figure 1

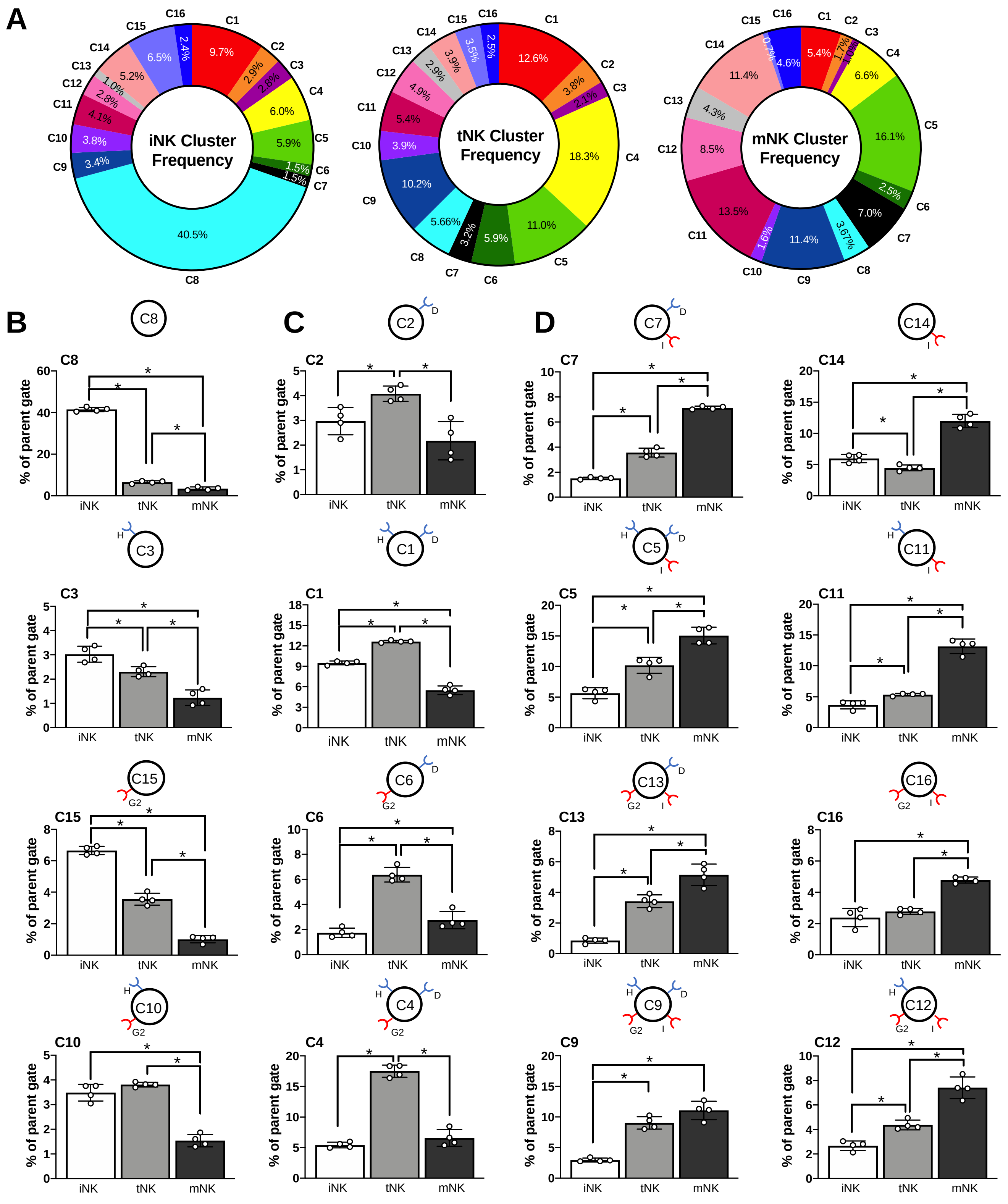


**Supplemental Figure 1. NK cell Ly49 cluster frequencies predominantly found at iNK, tNK and mNK cell stage. (A)** Summarized NK cell cluster mean frequencies gated on iNK (CD27+CD11b-), tNK (CD27+CD11b+) and mNK (CD27-CD11b+) cells and plotted on as part of a whole; (**B-C**) NK cell frequencies gated on parent maturation stage and grouped in clusters found predominantly in the (**B**) iNK, (**C**) tNK, (**D**) and mNK compartments. Asterisks indicate statistically significant differences between means as determined by the Student *t* test. *p < 0.05.

### Millan et al. 2020. Supplemental Figure 2

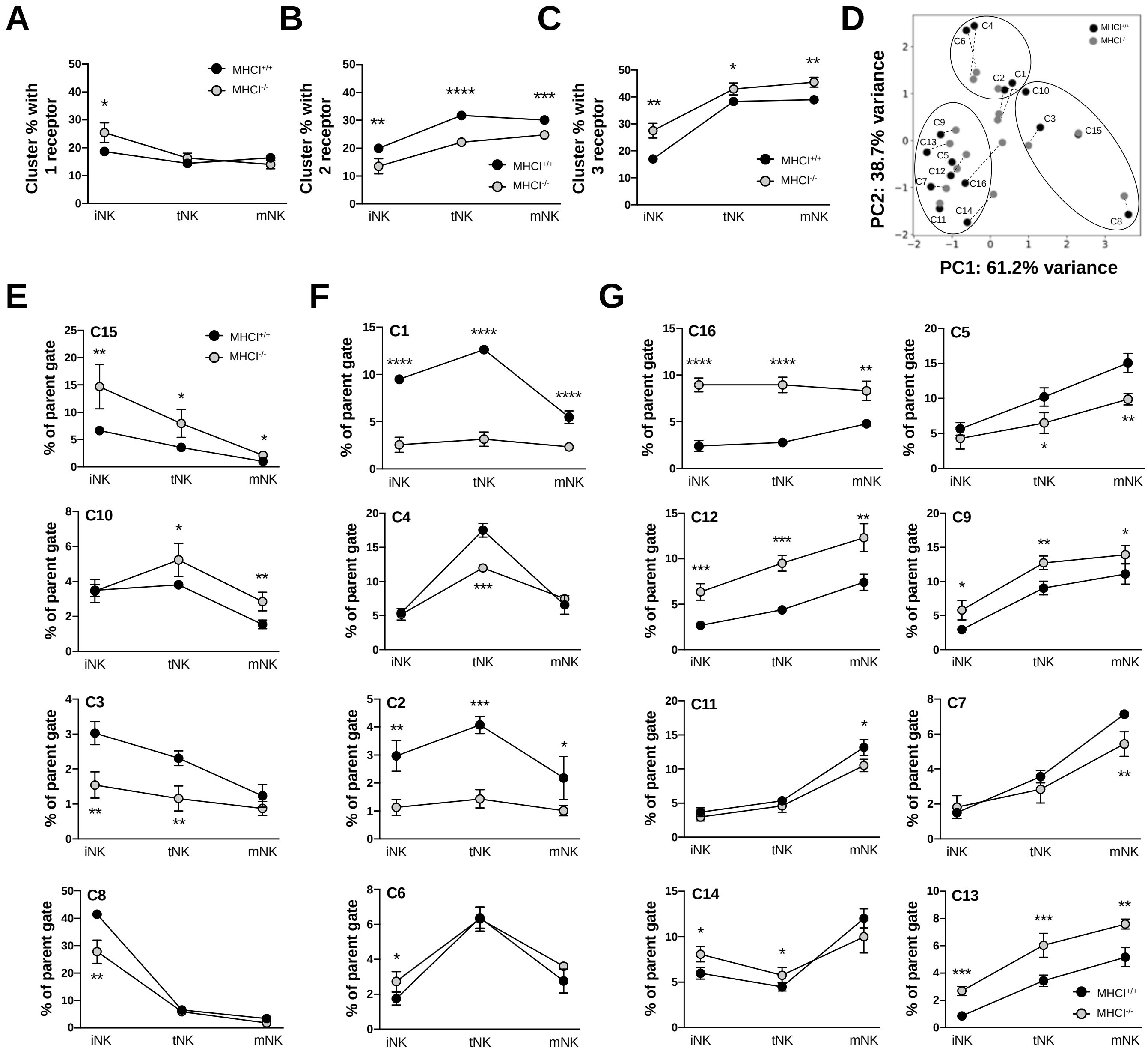


**Supplemental Figure 2. MHC-I$^{-/-}$ mice display altered frequencies of NK cell clusters at each maturation stage.** (**A**) Sum of frequencies of MHC-I$^{-/-}$ and MHC-I$^{+/+}$ NK cell clusters gated on iNK, tNK, and mNKs that express one, (**B**) two, and (**C**) three Ly49 surface receptors; (**D**) PCA analysis of calculated percent of NK cell clusters from a given iNK, tNK, or mNK stage of maturation between MHC$^{-/-}$ (grey dots) and MHC$^{+/+}$ (black dots) with a dotted line between the clusters to show changes between the two groups; (**E-G**) NK cell frequencies gated on parent maturation stage and grouped in clusters found predominantly in the (**E**) iNK, (**F**) tNK, (**G**) and mNK compartments for MHC-I$^{-/-}$ and MHC-I$^{+/+}$ mice. Asterisks indicate statistically significant differences between means as determined by the Student *t* test. *p < 0.05, **p < 0.01, ***p < 0.001, ****p < 0.0001.
